# Red blood cell dynamics in extravascular biological tissues modelled as canonical disordered porous media

**DOI:** 10.1101/2022.06.18.496666

**Authors:** Qi Zhou, Kerstin Schirrmann, Eleanor Doman, Qi Chen, Naval Singh, P. Ravi Selvaganapathy, Miguel O. Bernabeu, Oliver E. Jensen, Anne Juel, Igor L. Chernyavsky, Timm Krüger

**Affiliations:** School of Engineering, Institute for Multiscale Thermofluids, University of Edinburgh, UK; Manchester Centre for Nonlinear Dynamics, University of Manchester, UK; Department of Physics and Astronomy, University of Manchester, UK; Department of Mathematics, University of Manchester, UK; Department of Mechanical Engineering, School of Biomedical Engineering, McMaster University, Canada; Centre for Medical Informatics, Usher Institute, University of Edinburgh, UK; Bayes Centre, University of Edinburgh, UK; Maternal and Fetal Health Research Centre, School of Medical Sciences, University of Manchester, UK

**Keywords:** Haemodynamics, Red blood cells, Biological tissues, Porous media, Lattice-Boltzmann, Microfluidics

## Abstract

The dynamics of blood flow in the smallest vessels and passages of the human body, where the cellular character of blood becomes prominent, plays a dominant role in the transport and exchange of solutes. Recent studies have revealed that the micro-haemodynamics of a vascular network is underpinned by its interconnected structure, and certain structural alterations such as capillary dilation and blockage can substantially change blood flow patterns. However, for extravascular media with disordered microstructure (e.g., the porous intervillous space in the placenta), it remains unclear how the medium’s structure affects the haemodynamics. Here, we simulate cellular blood flow in simple models of canonical porous media representative of extravascular biological tissue, with corroborative microfluidic experiments performed for validation purposes. For the media considered here, we observe three main effects: first, the relative apparent viscosity of blood increases with the structural disorder of the medium; second, the presence of red blood cells (RBCs) dynamically alters the flow distribution in the medium; third, increased structural disorder of the medium can promote a more homogeneous distribution of RBCs. Our findings contribute to a better understanding of the cellscale haemodynamics that mediates the relationship linking the function of certain biological tissues to their microstructure.

## 1 Introduction

Significant progress in computational modelling has been made over recent years to elucidate the complex behaviour of blood flow in physiological environments, *e.g*., the small-vessel network in the brain [1, 2, 3], in the eye [4, 5], in tumours [6], and in microaneurysms [7]. However, the flow and transport of blood and solutes in other (*e.g*., extravascular) types of biological media with vital function, such as the intervillous space (IVS) of the human placenta featuring a highly disordered network of pores and flow passages of size comparable to that of red blood cells (RBCs), remain poorly understood [8, 9].

Comprehensive theories of flow and transport in porous media have been established, revealing subtle relationships between pore-scale structural heterogeneity and macroscopic flow properties [10, 11, 12, 13]. However, existing models of flow through porous media often assume a homogeneous fluid and cannot accurately infer the intricate blood rheology in living biological media, such as the IVS, where the particulate and highly-confined character of blood plays a key role, introducing spatiotemporal variability and nonlinearity beyond the description of prevalent continuum models [14, 15].

Emerging cell-resolved models of blood flow using advanced mesoscopic methods [16, 17, 18, 19] have been extensively applied since the late 2010s to simulate multiscale haemodynamics in synthetic or realistic vascular networks. These simulations have greatly improved our understanding of microscopic processes in the blood stream mediated by the flowing RBCs within, *e.g*., biased haematocrit distribution and oxygen transport arising from abnormal branching patterns of the vasculature [20].

Facilitated by robust image segmentation and meshing techniques [21, 22], cell-resolved models are now technically applicable to living porous media. These models provide a promising avenue for microscopic characterisation of the microstructure-dependent cellular blood flow, which can inform generalised constitutive relationships. A key aim is the development of effective reduced-order models for simulating large tissue/organ systems [23]. Additionally, cellular simulations can help design and optimise microfluidic oxygenators that serve as artificial lung assist devices [24].

In this work, we aim to characterise microscopic blood flow in canonical porous medium models constructed to represent simplistic extravascular biological media, for which we have control over the structural characteristics that can be related to physiological or pathological conditions of porous tissues and organs. Specifically, our focus is to quantify the correlations between cell-scale haemodynamics (e.g., flow patterns, RBC partitioning, haematocrit distribution) and key metrics of the porous medium, such as porosity and disorder. Primarily, we tackle the task computationally, with the aid of microfluidic experiments in equivalent flow systems for validation.

## 2 Methods

### 2.1 Design of canonical disordered porous media

We consider two typical designs of planar (quasi-2D) canonical disordered porous media (DPM) for our 3D simulations and validating experiments: *locally perturbed media (LPM)* (Fig. 1A) representing weak-disorder systems and *globally random media (GRM)* (Fig. 1C) representing strong-disorder systems, both of which are in the form of non-overlapping uniformly sized cylinders. The design process is inspired by the “bundle of tubes (uniform)” model implemented by Gostick *et al.* [25].

**Figure 1:**
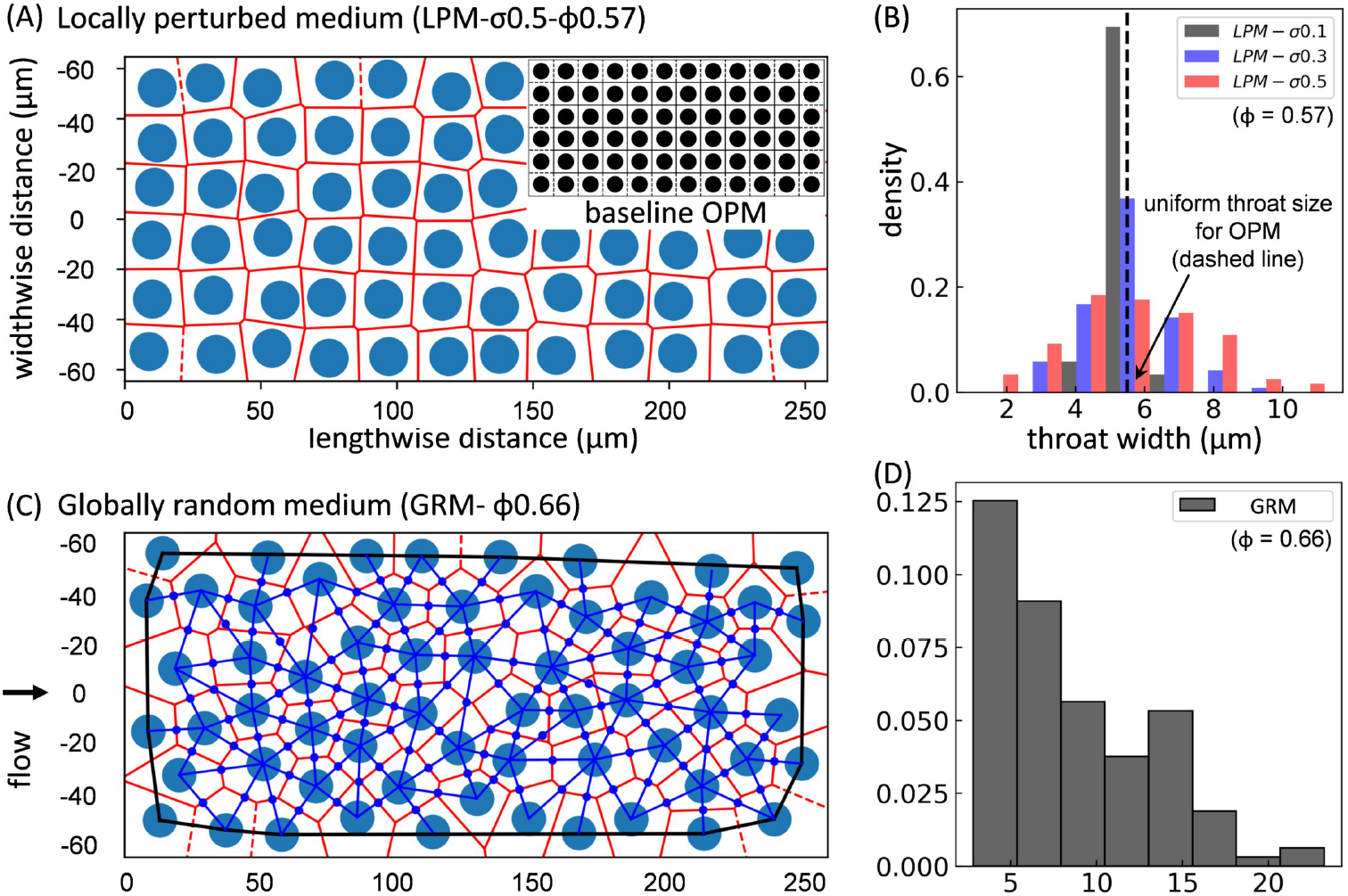
Construction of canonical porous media. (A) Locally perturbed medium (LPM), constructed by introducing local perturbations to a periodic ordered medium (OPM) of square obstacle arrays (inset). The throats (red edges) are delineated by Voronoi tessellation (red polygons); the pores are located where neighbouring edges meet. (B) Throat-width distributions (in the form of normalised histograms) for LPM geometries perturbed from an OPM of porosity *ϕ* = 0.57, with incremental disorder *σ* = 0.1, 0.3, 0.5. (C) Globally random medium (GRM) with pores and throats (excluding those in direct contact with the boundaries, which are located outside the black bounding box) delineated by Voronoi tessellation (red polygons) and Delaunay triangulation (blue triangles). The blue dots indicate the middle points of pore openings where local flows and cell fluxes are evaluated. (D) Throat-width distribution for GRM (*ϕ* = 0.66), same axis labels as (B). Flow is driven in the horizontal direction.

To obtain LPM, we first construct an ordered porous medium (OPM) by placing cylinders of constant diameter *D_c_* (determined by porosity *ϕ* and domain size) on a square grid with length *L* and width *W* (Fig. 1A inset), which can be regarded as a generalised capillary bundle model. Three porosity values *ϕ* = 0.57, 0.62, 0.67 are considered, which lead to a confinement ratio (defined as *χ* = D_RBC_/W_throat_, denoting the ratio of effective RBC diameter to throat width) in the baseline OPM geometry of *χ* = 1.2, 1.0, 0.9, respectively. The alteration of porosity is achieved by varying *D*_c_ while keeping the cylinder positions unchanged. The structural disorder of the LPM is then created by introducing random perturbations to the cylinder positions (*x*,*y*) with standard deviation *σ* = 0.1, 0.3, 0.5, 0.7, where (*x,y*) are drawn from a bivariate normal distribution. For a relatively large *σ* (*e.g*., *σ* = 0.7), the perturbations are likely to cause overlapping cylinders and several attempts may be needed to construct a non-overlapping layout. The resulting distributions of throat widths in the LPM are Gaussian (Fig. 1B).

The GRM is constructed by placing a uniformly random spatial distribution of cylinders in a domain of given size for a designated porosity (*ϕ* = 0.66), while enforcing a minimal separation distance of 0.4 *D*_RBc_ between neighbouring cylinders (Fig 1C). The resulting distribution of throat widths in the GRM is approximately exponential (Fig 1D).

For ease of description when comparing various geometries in plots, each LPM geometry is assigned with a disorder identifier and a porosity identifier, *e.g.*, LPM-*σ*0.5-*ϕ*0.57 for LPM with *σ* = 0.5, *ϕ* = 0.57.

For OPM and GRM, only a porosity identifier is assigned, *e.g*., OPM-*ϕ*0.57 and GRM-*ϕ*0.66.

### 2.2 Numerical model

The immersed-boundary-lattice-Boltzmann method (IB-LBM [26]) simulates cellular blood flow as a suspension of deformable RBCs (see Sec. S1 in the **Supplementary Material** for more details of the model) through a shallow Hele-Shaw bed (*L* × *W* × *H* = 258 × 129 × 6 *μ*m^3^) of vertical cylinders, which is constructed through extrusion over a distance (*H*) of the desired DPM designs of dimension *L* × *W* (LPM or GRM) in the depth direction. Each RBC is modelled through the finite element method as a closed hyperelastic membrane with the unstressed shape of a biconcave discoid. The mechanical properties of the RBC membrane are controlled by elastic moduli accounting for different energy contributions, including the strain modulus *κ_s_*, bending modulus *κ_b_*, dilation modulus *κ_α_*, surface modulus and volume modulus *κ_V_* [27] (see Table S1). To tackle close cell-cell and cell-wall interactions, a repulsion potential decaying with inverse distance between neighbouring surfaces is numerically implemented with interaction intensities comparable to the bending elasticity of the RBC membrane [28].

In the simulations, an RBC-free plasma flow is initialised from left to right along the channel axis (length direction) with a designated volumetric flow rate *Q_0_* (Fig. 2A). The flow is driven by imposing a parabolic velocity profile at the inlet (assuming Poiseuille flow under *Q_0_*) and a reference pressure at the outlet (*p*_0_), through two cylindrical flow extensions smoothly stitched to the porous bed of rectangular cross-section (*W* × *H*). The no-slip condition is imposed at the wall and all fluid-solid interfaces. Once the plasma flow is converged, RBCs are randomly inserted from the inlet flow extension in a continuous manner with fixed RBC volume fraction (*i.e.,* feeding discharge haematocrit, *H_F_* = 0.1, 0.2, 0.3). RBCs reaching the outlet flow extension are removed from the system (see **Supplementary Movie1** for an example simulation in the OPM-*ϕ*0.57 geometry).

**Figure 2:**
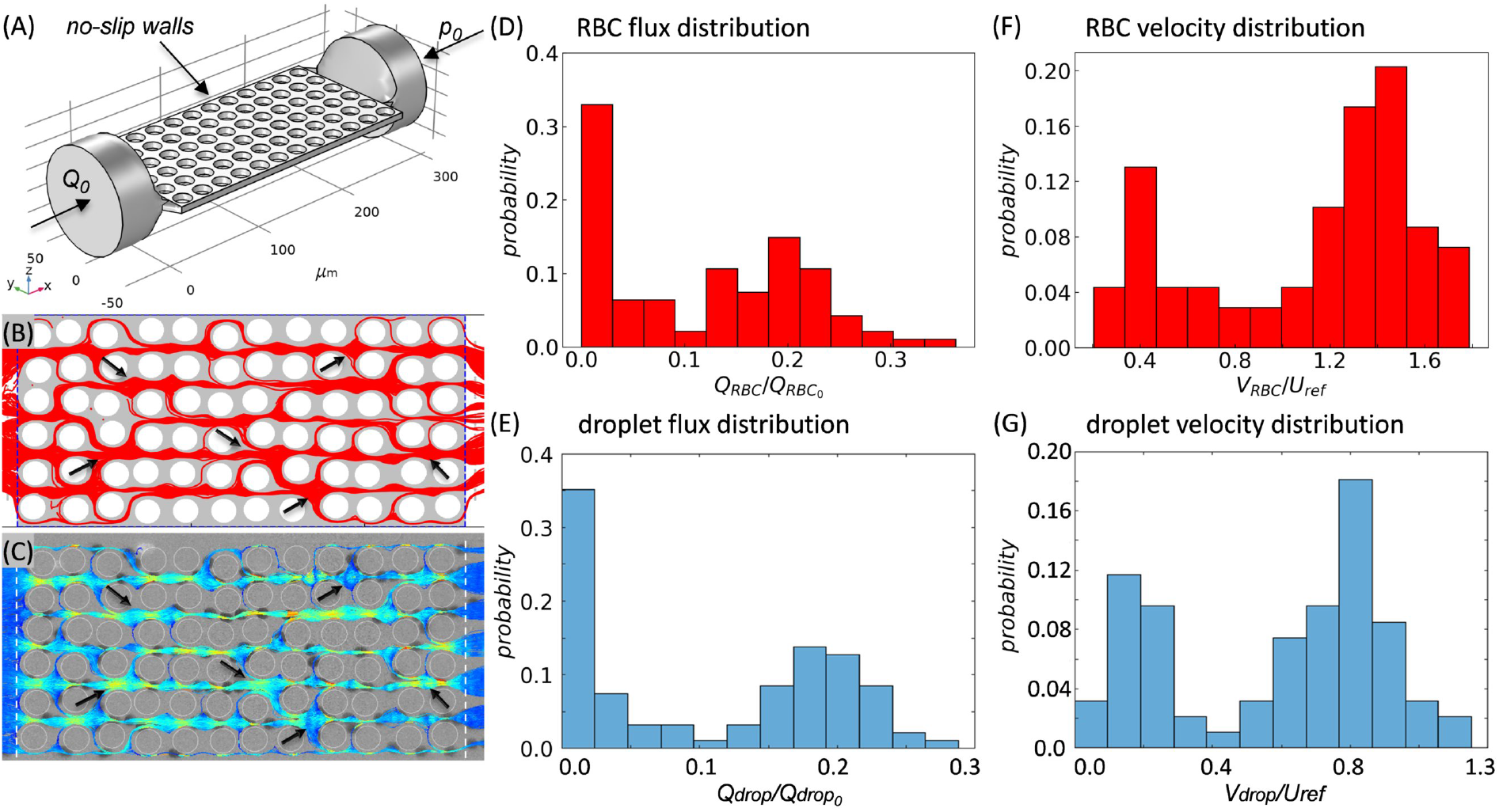
Cross-validation of simulation and experiment (LPM-*σ*0.5-*ϕ*0.57). (A) Schematic of simulation domain and boundary conditions. (B,D,F) Simulation results. (C,E,G) Experiment results. (B,C) Superimposed particle trajectories over time. The red lines in (B) represent the RBC trajectories, and the red dots indicate occluded throats by the RBCs. The coloured lines in (C) represent the droplet trajectories with high-velocities marked in brighter colours. The black arrows in (B,C) highlight example regions of interest where simulation and experiment agree well. (D,E) Distributions of normalised particle fluxes. (F,G) Distributions of normalised particle speed.

### 2.3 Microfluidic experiments

In the microfluidic experiments, a suspension of silicone oil droplets (Sigma Aldrich, viscosity *ν* = 20 cSt with paraffin oil dye) in a mixture of water and glycerol (Sigma Aldrich, 80:20 by volume with 0.2% SDS, viscosity 0.24 Pa s at 21°C) is used as biomimetic model for blood. The droplets were generated in a flow-focusing device [29] which was connected to the porous medium via a bent glass capillary (length 100 mm, outer diameter 1 mm, inner diameter 0.58 mm). Droplets with a diameter of 240 *μ*m were produced by flowing the inner phase (silicone oil) at 4 *μ*L/min and introducing the outer phase (water/glycerol) through a cross-junction at 5 *μ*L/min immediately upstream of the flow-focusing channel constriction. Downstream of the droplet formation, another 11 *μ*l/min of water-glycerol mix was added to set the droplet flow fraction to *H_F_* = 0.2. Flow control at the different inlets was achieved with a combination of a pressure controller (Elvesys) fitted with flow meters and a syringe pump (KD Scientific). Both the flow-focusing device and the porous medium were made from PDMS (Polydimethylsiloxane, Sylgard 184, Dow Corning). To obtain hydrophilic wetting behaviour, the devices were oxidised by oxygen plasma treatment (Henniker plasma HPT-100) and immediately filled with water. The porous medium corresponds to the LPM-*σ*0.5-*ϕ*0.57 geometry (i.e., disorder 0.5 and porosity 0.57) used in numerical simulations, scaled by the ratio of the droplet diameter to the equivalent diameter of simulated RBCs (*D*_RBC_ = 6.68 μm). The suspension flow was imaged with a monochrome CMOS camera (PCO 1200hs) and analysed off-line using the ImageJ platform TrackMate [30].

### 2.4 Cross-validation of simulation and experiment

Good agreement between the simulated and experimented particle dynamics is obtained (see Fig. 2B–G and **Supplementary Movie2–Movie3** for comparison in an example geometry), including the pattern of particle trajectories (Fig. 2B,C), the distribution of particle fluxes (Fig. 2D,E) and the distribution of particle velocities (time-average speed for particles crossing a given pore-throat, Fig. 2F,G). Both the RBCs and droplets demonstrate evident preferential pathways or regional shunts when travelling across the porous domain, with discrete clusters of high-flux and low-flux throats. Similarly, the mean particle velocities at individual throats feature a bimodal distribution where high-velocity and low-velocity populations co-exist.

Here, particle fluxes *Q*_RBC_ (or *Q*_drop_) and particle speed *V*_RBC_ (or *V*_drop_) are evaluated at individual throats in the porous domain. Taking the simulation for example, *Q*_RBC_ is defined as

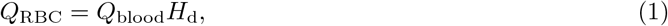

where *Q*_blood_ is the local volumetric flow rate (denoted as *Q*_p_ in plasma flow and *Q*_s_ in suspension flow) across a given throat and *H*_d_ is the discharge haematocrit measured for the same throat. The normalising particle flux *Q*_RBC0_ is chosen as the overall particle flux fed at the flow inlet; it is defined as

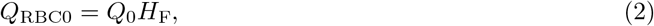

where *Q*_0_ is the flow rate imposed at the inlet. The normalising particle velocity *U*_ref_ is chosen as the peak fluid velocity evaluated at the centre of horizontal pores in the baseline OPM geometry, defined as

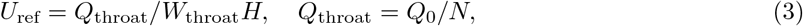

where the throat width *W*_throat_ measures the minimum distance between two neighbouring cylinders and *N* is the number of throats across the vertical direction (perpendicular to flow axis) in the OPM geometry. Note that *U*_ref_ should not be mistaken for *U*_0_, which is the superficial velocity *U*_0_ defined as

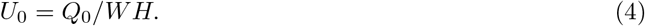

*U*_0_ is adopted as spatially averaged advection speed to estimate the reference transit time *T*_ref_ for particles to travel across the whole domain length, defined as

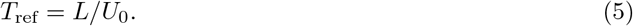

## 3 Results and Discussion

We first demonstrate the effect of porous medium structure and the presence of RBCs on the flow resistance. Then, we investigate the RBCs’ contribution to pore-scale flow redistribution and the RBCs’ spatiotemporal dynamics in relation to structural characteristics of the porous domain.

### 3.1 Effect of structural disorder on flow resistance in porous media is more prominent with RBCs present

To assess the resistance to plasma flow (RBC-free) and suspension flow (RBC-laden, with default feeding haematocrit *H*_F_ = 0.2), we compare the overall pressure drop Δ*P* across different porous media under an identical feeding volumetric flow rate at the inlet (*Q*_0_ = 0.4 μL/min in simulations, unless otherwise specified). For suspension flow, we also examine the relative apparent viscosity *η*_rel_ comparing the suspension apparent viscosity *η*_app_ to that of plasma *η*_0_ (1×10^-3^ Pa s), which can be calculated as the ratio of pressure drops for equivalent suspension and plasma flows under same inflow (*Q*_s_ = *Q*_P_ = *Q*_0_), *i.e*., *η*_rel_ = *η*_app_/*η*_0_ = ΔP_s_/ΔP_p_.

The porosity is found to have a prominent effect on the plasma flow resistance (Fig. 3A). For instance, a decrease of *ϕ* = 0.67 to 0.62 and 0.57 for the intermediately disordered LPM (*σ* = 0.5) leads to 25.4% and 65.1% rise in the pressure drop, respectively. Similarly, the suspension also experiences higher flow resistance: 25.6% and 66.7% increase, respectively (Fig. 3A). However, the relative apparent viscosity of the suspension remains approximately constant, therefore suggesting a small effect of porosity within the studied range (Fig. 3A).

**Figure 3:**
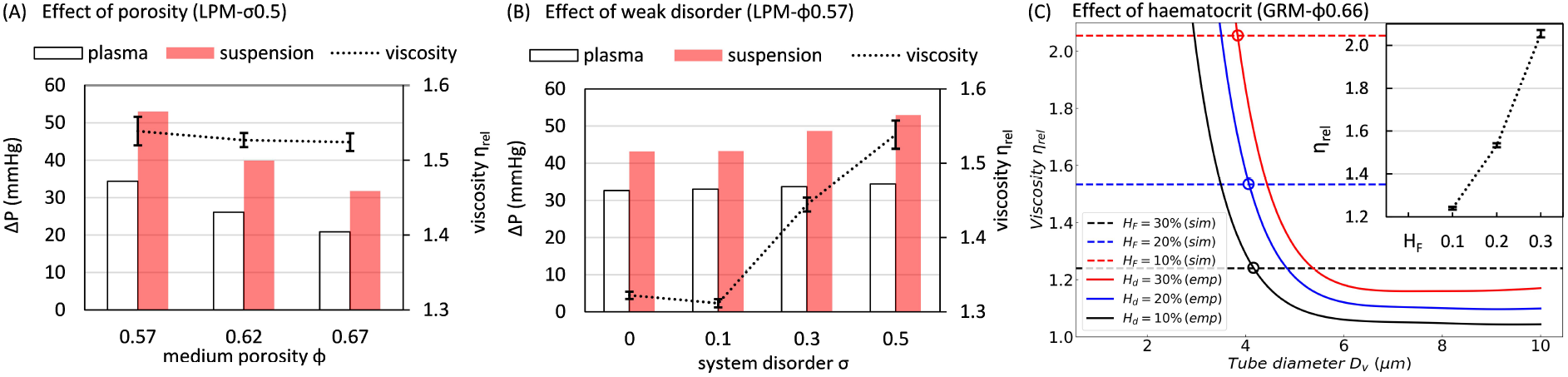
Evaluation of flow resistance in porous media with different geometries and flow conditions. Pressure drop across the simulated porous media under designated volumetric flow rates *Q*_0_ fed at the inlet (*Q*_0_ = 0.4μL/min unless specified otherwise). The default porosity is *ϕ* = 0.57. (A) Variation of pressure drop against porosity in LPMs of fixed disorder *σ* = 0.5 for plasma flow and a suspension flow with RBCs of feeding haematocrit *H*_F_ = 0.2. The relative apparent viscosity *η*_rel_ of the suspension flowing within is assessed through comparing with plasma viscosity. The error bars represent standard deviation of measurements from five time instants of the suspension flow. (B) Variation of pressure drop against disorder (*σ* = 0.1, 0.3, 0.5) in LPMs of fixed porosity *ϕ* = 0.57 for plasma flow and a suspension flow with *H*_F_ = 0.2, where *η*_rel_ is quantified the same way as in (A). (C) Comparison of *η*_rel_ (dashed horizontal lines) in the simulated porous media with empirical predictions (solid curved lines) for straight tubes *in vitro* [31]. The round symbols indicate the characteristic diameter *D*_v_ of an equivalent tube within which the suspension has the same relative apparent viscosity. The inset of (C) shows the variation of simulated *η*_rel_ against increasing *H*_F_ =0.1, 0.2, 0.3 in the GRM (*ϕ* = 0.66).

For a given porosity of *ϕ* = 0.57, our simulations in LPM geometries reveal a marginal effect of incremental disorder on the resistance of plasma flow, but an evident effect on that of the suspension flow (Fig. 3B). The relative apparent viscosity increases substantially beyond a critical degree of disorder; *η*_rel_ is 16.4% higher at *σ* = 0.5 compared to its value in the ordered medium (*σ* = 0).

We further investigate the effect of stronger disorder on the relative apparent viscosity of suspension flow with the GRM geometry where random disorder is introduced globally (Fig. 3C). Different haematocrit levels are examined, with the RBC volume fraction varying between *H*_F_ = 0.1 – 0.3, which brings up the relative apparent viscosity by nearly 66%. The viscosity values (*η*_rel_ ∈ [1.24, 2.05]) are in line with recent experimental results for a large-scale porous domain consisting of hexagonal arrays of circular pillars separated by a constant distance of 10 *μ*m [32]. In our case, this distance is represented by the throat width *W*_throat_. For the GRM geometry *W*_throat_ is not constant, but rather has an exponential distribution with a mean value of 8.5 *μ*m and a median of 7.1 *μ*m (*W*_throat_ ∈ [2.8, 23.3] *μ*m). Through comparison with an established empirical model for blood viscosity *in vitro* [31] (see the formulation in Sec. S2 of the **Supplementary Material**), our simulated suspension flow at *H*_F_ = 0.1, 0.2, 0.3 through the GRM domain is found to have equivalent relative apparent viscosity to a uniform cylindrical tube of 4.2 *μ*m, 4.1 *μ*m, 3.8 *μ*m, respectively (Fig. 3C).

The significantly different effects of porosity and disorder on the relative apparent viscosity is surprising; both decreasing *ϕ* and increasing *σ* are expected to contribute to stronger pore-scale confinement that impedes the transport of RBCs [33] (see definition of confinement ratio *χ* in Sec. 2.1). We believe this difference in behaviour stems from the sensitivity of local confinement to changes in *ϕ* and *σ*. Changing *ϕ* uniformly modifies all throats (for *ϕ* ∈ [0.57, 0.67], the mean throat width only varies in the range of 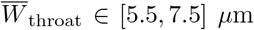). In contrast, a change in *σ* augments the possibility of severe confinement in the system by broadening the *W*_throat_ distribution and giving rise to exceedingly small throats (*W*_throat_ < 3 *μ*m, see Fig. 1B). Note, however, that the impact of occasional severe confinement in some throats when increasing *σ* might be partially compensated by the simultaneously introduced larger throats, leading to an overall minor increment in the relative apparent viscosity.

Theoretically, when the size of a channel approaches that of an RBC, the cell-free layer vanishes due to confinement and the cells are in direct contact with the wall, consequently increasing the relative apparent viscosity of blood in straight tubes as postulated by the well-known Fåhræus-Lindqvist effect [31] (Fig. 3C). However, depending on its severity, such confinement may also contribute to lower relative apparent viscosity by forcing the organisation of RBCs into streamlined single-file form, thus excluding excessive cell-cell collisions that would otherwise increase flow resistance [14]. For the disordered geometries studied here, where a tightly interconnected network of pores and throats (rather than isolated channels) are involved, it is expected that both effects exist and compete with each other in determining the relative apparent viscosity of the suspension flow. Further, because each throat has a varying crosssection of contraction and expansion, the spatial organisation of RBCs constantly re-arranges without reaching a fully developed profile, which constitutes a mechanism for higher relative apparent viscosity in the porous medium than straight tubes of geometrically equivalent diameter (Fig. 3C). These counteracting mechanisms associated with structural alterations sometimes balance each other and the overall viscosity of the suspension remains roughly unchanged under certain circumstances *(e.g.,* Fig. 3A).

### 3.2 RBC traffic dynamically alters the pore-scale flow distribution

For both the plasma flow and suspension flow (Fig. 4A), a larger degree of structural disorder (i.e., increasing *σ*) is found to promote higher perfusion of the pore space network as indicated by a smaller percentage of throats experiencing negligible flux (compare Fig. 4B with Fig. S1A in the **Supplementary Material**). In both RBC-free and RBC-laden flow scenarios, the flow distributions evolve from discrete high-/low-flow clusters towards a continuous exponential-like profile as the level of medium disorder increases, similar to earlier findings on Newtonian flows through perturbed cylinder arrays [11]. However, this evolution is noticeably accelerated in the presence of RBCs, *i.e*., achieving an exponential-like flow distribution at lower level of disorder (Fig. 4B). Given that such exponential distribution is underpinned by less extreme local flow fractions at nodal points (*i.e.*, bifurcations) [11], the RBCs contribute to more robust perfusion of the porous domain, which is in line with a recent report on the role of RBCs in stabilising blood flow in the capillary bed [34]. Indeed, for strongly disordered geometry where flow channelisation (known to be associated with anomalous solute transport in heterogeneous media [35]) prevails in the plasma flow, the presence of RBCs is found to break down some dominant pathways and enhance regional perfusion (see Fig. S1B in the **Supplementary Material**).

**Figure 4:**
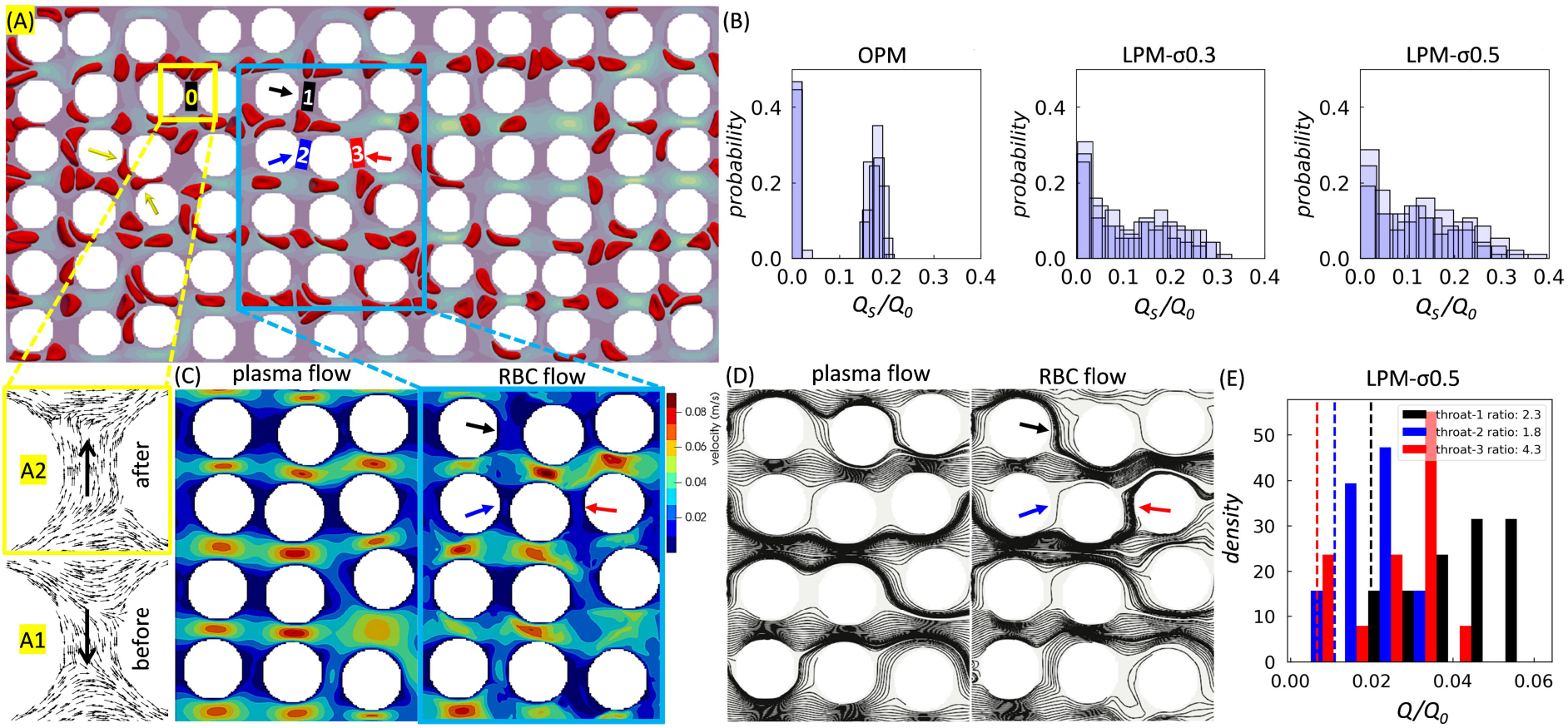
Effect of RBC traffic on local flow patterns. The porosity is *ϕ* = 0.57, and the feeding haematocrit is *H*_F_ = 0.2. Flow is from left to right. (A) Snapshot of an LPM simulation (*σ* = 0.5). The arrows (yellow) near the inlet indicate two cell-occlusion events. A small rectangle (yellow) marks a throat nearby (“0”) within which flow reversal can be observed before (inset “A1”) and after (inset “A2”) the cell-occlusion events. Another rectangle (blue) marks the central region enlarged in (C,D) for pore-scale details. (B) Flow rate distributions for RBC flows through LPM with increasing levels of disorder *σ* = 0, 0.3, 0.5 (where *σ* = 0 refers to the OPM). The histograms show flow rate magnitudes *Q*_s_ evaluated at all throats within the pore space network, and *Q*_0_ = 0.4 *μ*L/min is the imposed flow rate at the inlet. Three time instants are superimposed to show the temporal fluctuations due to the dynamics of discrete RBCs. (C) Instantaneous flow field (plasma-only and suspension) for the same *Q*_0_ = 0.4*μ*L/min (*σ* = 0.5). (D) Instantaneous streamlines for the same scenarios as in (C). The coloured arrows in (C,D) point to the three throats of interest (“1”, “2”, “3”) marked in panel (A). (E) Quantification of the time-dependent RBC flow through the designated throats of interest as in (C,D). The histogram for each throat shows the probability of finding the instantaneous flow rate magnitude *Q* normalised by the imposed inflow *Q*_0_; the histograms are obtained by evaluating fifteen time instants. The vertical dashed lines represent the corresponding steady and normalised plasma flow rates (magnitude) *Q*_P_/*Q*_0_ (note that the two lines for throats “1” and “3” have been moved away from each other by a finite distance for visual clarity, which would otherwise overlap). The ratios in the legend indicate the median suspension flow rate divided by the constant plasma flow rate in each throat investigated.

Examination of the pore-scale dynamics reveals that the presence of RBCs introduces intermittent and/or permanent occlusion of narrow throats (Fig. 4A and **Supplementary Movie2**), featuring trapped RBCs between adjacent cylinders as observed in a recent experimental study [36]. Accompanied by such occlusion events are occasional flow reversals found in throats located in the vicinity (see Fig. 4A insets, “A1” and “A2”). These occlusions may in turn contribute to the diversion of flow *(via* occlusive pressure feedbacks [37]) to otherwise poorly infused regions (*e.g.*, vertically-oriented throats) and lead to more evenly distributed suspension flow in the system (see the enhanced transverse flows in Fig. 4C, right panel) compared with plasma-only flow (Fig. 4C, left panel) under identical inflow conditions. Indeed, the underlying streamline patterns in Fig. 4D suggest that some negligibly perfused throats in the plasma scenario become better perfused in the suspension scenario.

Furthermore, the RBCs impose a temporal signature on the pore-scale flow which is characterised by distinct patterns of flow channelisation over time (Fig. S2 in the **Supplementary Material**). To quantify the temporal variation, three throats of interest (Fig. 4E) are analysed for their instantaneous flow rates over fifteen time instants. Wide distributions of flow rates are found, and the time-averaged blood flow at the throats is roughly two to four times as high as in the plasma-only scenario. It is noteworthy that despite the highly dynamic RBC traffic and flow fluctuations at the pore scale, the pressure drop across the entire porous bed is essentially steady (see Fig. 3 for the narrow error bar ranges).

### 3.3 Increased disorder promotes homogeneous RBC transport and haematocrit distribution

RBC trajectories in the porous bed provide information about the spatiotemporal dynamics of the cells (compiled with over 1000 cells, Fig. 5A). Fig. 5B shows the corresponding distributions of RBC transit times in geometries with different medium disorder. The data reveal that both the shape of the distribution and the median transit time are quantitatively similar in cases investigated here (*ϕ* = 0.67), provided the porosity remains constant (confirmed for cases under *ϕ* = 0.57 in Fig. S3A of the **Supplementary Material**). In other words, the transit time distributions are insensitive to the medium disorder under our studied range of porosity magnitudes. The insensitivity is likely caused by two opposing effects of structural disorder on the motion of RBCs: increasing the tortuosity of existing pathways slows down individual cell transits, but enables more flow pathways by creating local transverse pressure gradients. Our results suggest that, in a disordered porous medium such as the placental IVS [22], the average residence time of RBCs (associated with oxygenation and solute transport processes) within representative tissue volumes can be independent of the inherent structural heterogeneity if the tissue porosity remains roughly the same.

**Figure 5:**
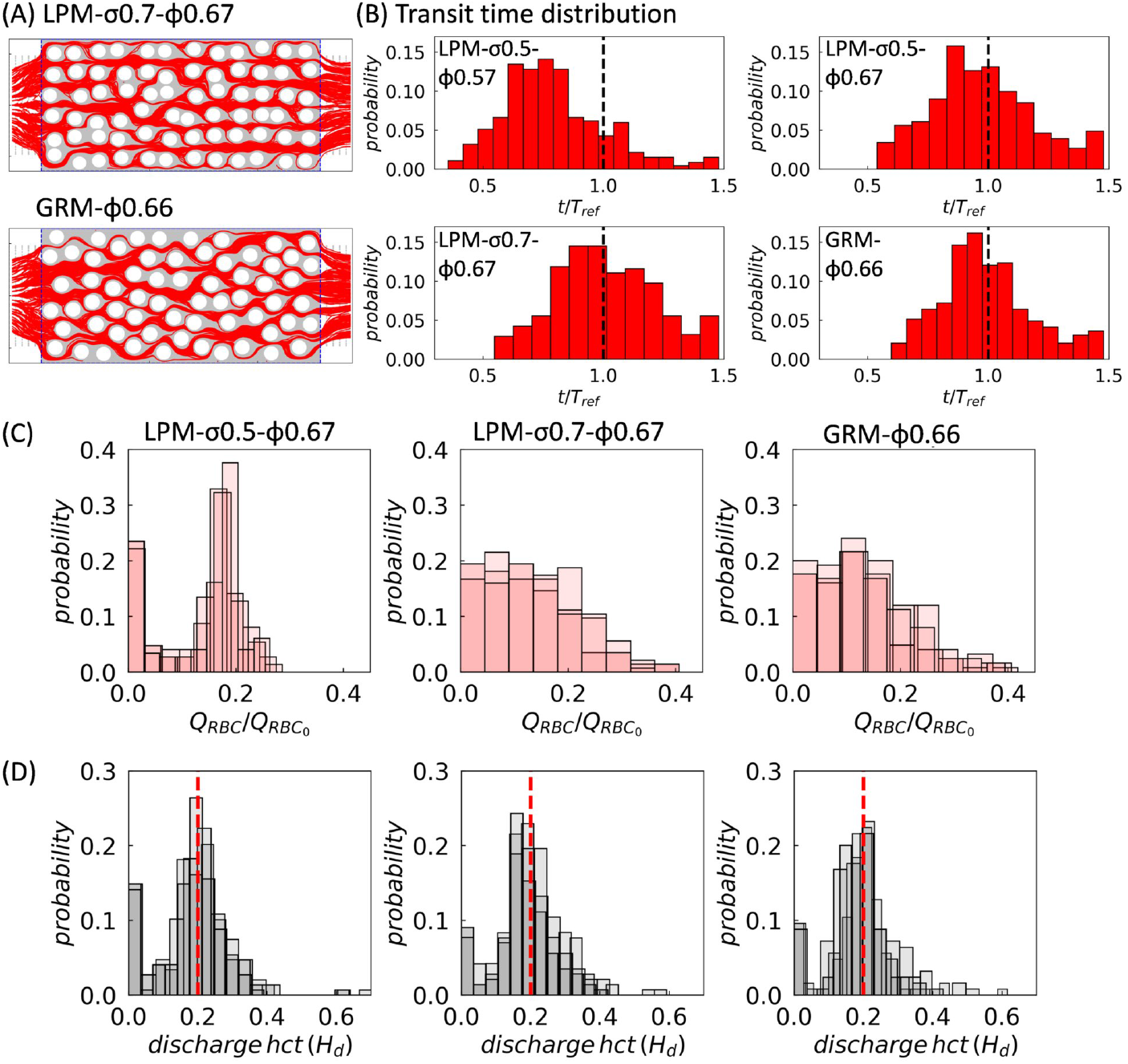
Effect of disorder on RBC perfusion and haematocrit distribution. (A) Combined RBC trajectories in simulation over a time period of 45 ms for LPM-*σ*0.7-*ϕ*0.67 and GRM-*ϕ*0.66. (B) Distributions of normalised RBC transit time subject to varying levels of porosity and disorder. The transit time is calculated based on each RBC entering and leaving the porous domain between the two vertical dashed lines (blue) in (A). The reference transit time *T*_ref_ was defined in Sec. 2.4. (C) RBC flux distributions corresponding to the high-porosity (*ϕ* = 0.66-0.67) geometries in (B). The superimposed histograms from three consecutive time intervals for each geometry show the distribution of RBC fluxes evaluated at individual throats throughout the pore space network at three consecutive time intervals. (D) Distributions of discharge haematocrit *H*_d_ corresponding to the simulated RBC fluxes in (C).

On the other hand, increasing the porosity from *ϕ* = 0.57 to *ϕ* = 0.67, while keeping the disorder unchanged, leads to an *increase* in median transit time (compare “LPM-*σ*0.5-*ϕ*0.57” and “LPM-σ0.5-*ϕ*0.67” in Fig. 5B). This seemingly counter-intuitive effect (*i.e.,* slower flow under relaxed confinement) is due to an overall smaller interstitial flow speed at higher porosity under the imposition of a *constant* feeding flow rate *Q*_0_ (rather than constant pressure drop Δ*p*). The evidently altered pattern of RBC transit times by a moderate change in the medium porosity suggests a potential mechanism for impaired placental function in pre-eclamptic or diabetic pathologies, where the porous tissues have been found either overly sparser or denser than under normal conditions [38].

Finally, we examine the spatiotemporal RBC distribution in more detail. A visual inspection of the time-lapse RBC perfusion reveals that there are still plenty of throats devoid of RBCs for the intermediate disorder *σ* = 0.5 (**Supplementary Movie4**); however, for higher degree of perturbed disorder *σ* = 0.7 and the random disorder (Figure 5A and **Supplementary Movie5–6**), the pore space network is more strongly perfused (also see). Fig. 5C shows the RBC flux distributions in the throats of the networks. These data confirm that the locally-perturbed “LPM-σ0.7” and the globally-random “GRM” have comparable RBC distributions while the less disordered “LPM-*σ*0.5” features a large number of throats without RBCs.

The discharge haematocrits *H*_d_ of individual throats tend to be close to the systemic haematocrit *H*_F_ = 0.2 as the structural disorder becomes sufficiently large in the system *(e.g*., *σ* ≥ 0.5), featuring a primary peak located around *H*_d_ = *H*_F_ in the distribution profile (Fig. 5D). This concentrated distribution around *H*_d_ = 0.2 is still visible for weakly disordered geometries with *σ* ≤ 0.3, but outweighed by a more dominant peak at zero (Fig. S3B in the **Supplementary Material**).

## 4 Conclusion

We have investigated the micro-haemodynamics of cellular blood flow in a class of porous media as a canonical model for extravascular geometries representative of biological tissues such as the porous intervillous space in the human placenta. Two types of canonical porous media with varying degrees of porosity were considered: weakly disordered porous media based on a regular square array of circular obstacles in a flat channel, and a random porous medium with non-overlapping circular obstacles. We employed simulations combining the lattice-Boltzmann, finite-element and immersed-boundary methods to explore the effects of haematocrit, porosity and structural disorder on the blood rheology, perfusion and distribution of red blood cells (RBCs) in the porous media. We used scaled-up experiments with droplets to validate the simulation framework. We found that the flux and velocity distribution of the simulated RBCs and measured droplets are qualitatively and quantitatively similar.

The findings from the simulations are threefold:

1. The relative apparent viscosity of cellular blood increases with the structural disorder of the porous media considered here, but it is largely independent of the studied porosity. These counter-intuitive findings are likely caused by competing effects due to reduction of the cell-free layer thickness and cell ordering in narrow channels.
2. The presence of RBCs dynamically alters the flow distribution in the porous media investigated here. Throats that are only weakly perfused in the absence of RBCs can receive significantly higher flow due to RBCs partially blocking the flow in other, better-perfused throats. Due to the motion of the RBCs, the flow rates in the throats can fluctuate strongly.
3. Increasing the structural disorder of the porous media considered here promotes a more homogeneous distribution of RBCs as measured by cell fluxes and discharge haematocrits. While RBCs favour fast lanes through ordered or weakly disordered porous media, a stronger perturbation creates new pathways for the RBCs, leading to a more homogeneous haematocrit distribution throughout the geometry.

We have presented an experimental-computational framework for studying the micro-haemodynamics in disordered porous media. Our framework can be extended to more realistic architectures of biological tissues and organs. We envision that a multi-disciplinary approach, cross-validating simulations and experiments to extract generalised constitutive relationships for RBC flow in complex geometries, will help bridge the gap between microscopic characterisation and tissue/organ-level modelling which are both necessary to reveal the relationship between structure and function of biological tissue and organs.

## 5 Data accessibility

Data and information supporting this article are provided in the electronic supplementary materials. The source code for the blood flow simulations is available at https://github.com/hemelb-codes/hemelb.

## 6 Authors’ contributions

Q.Z. designed the study, analysed the data and wrote the manuscript; Q.Z. (computational modelling), K.S., Q.C. and N.S. (microfluidic experiments), and E.D. (mathematical analysis) performed the research; P.R.S., M.O.B. and O.E.J. discussed the results and revised the manuscript; A.J., I.L.C. and T.K. supervised the study and edited the manuscript. All authors critically reviewed the manuscript and approved the final version for publication.

## 9 Acknowledgements

The authors acknowledge the computational support provided by Rupert W. Nash (EPCC, The University of Edinburgh) and the *HemeLB* development team to this work.

## Supplementary Materials

### S1 Numerical model for cellular flow simulation

The immersed-boundary-lattice-Boltzmann method (IB-LBM [27]) is employed to simulate cellular blood flow as a suspension of deformable RBCs. The numerical model is implemented in the parallel flow simulator *HemeLB* [39] (open source at https://github.com/hemelb-codes/hemelb), which has been successfully applied to cellular simulation in complex and sparse geometries (*e.g.*, realistic microvasculature [4]). The 3D fluid flow governed by the Navier-Stokes equations is solved by the lattice-Boltzmann method (LBM) with standard D3Q19 lattice [40], BGK collision operator [41], Guo’s forcing scheme [42] and the Bouzidi-Firdaouss-Lallemand (BFL) implementation of no-slip condition at the walls [43]. Open boundary conditions for flow inlets/outlets are implemented with the Ladd velocity boundary condition [44]. To configure the simulation, each porous medium geometry (generated in STL format) is discretised into a uniform mesh of cubic voxels. The voxel size *(i.e.,* the lattice constant Δ*x*) is chosen such that the plasma flow can be reliably solved at the narrowest pore-throats in the simulated porous media. The relaxation time parameter *τ*_BGK_ is determined based on a previous study [45] for minimising relative error of simulated flow rates. Refer to Table S1 for values of key simulation parameters.

The discrete RBCs are modelled as triangulated Lagrangian membranes using the finite element method (FEM) and are coupled with the fluid flow using the immersed-boundary method (IBM [46]). Each RBC consists of N triangular facets and present a discocyte shape at rest. The mesh resolution of the membrane is chosen such that the average edge length of the facets matches the voxel size Δ*x* of the flow domain for numerical stability and accuracy [26]. The membrane is hyperelastic, isotropic and homogeneous, with its mechanical properties controlled by (*κ*_s_, *κ*_b_, *κ_α_*, *κ*_A_, *κ*_V_) as mentioned in the main text. *κ*_cc_ and *κ*_cw_ are intensities of the repulsive potential (decaying with inverse distance) between neighbouring cells, and neighbouring cell-wall, respectively. The cytosol enclosed by the RBC membrane is regarded as a Newtonian fluid with same viscosity to the plasma. The membrane viscosity is not considered in the present material model. Refer to Table S1 for values of the above parameters.

**Table S1:** Key parameters of the flow and RBC model. “~” means dimensionless simulation units.

| Parameter | Description | Value | Unit | Comment |
| --- | --- | --- | --- | --- |
| $\Delta x$ | lattice constant | 0.4 | $\mu\text{m}$ | decided by minimal throat widths |
| $\Delta t$ | time step length | $1.6 \times 10^{-8}$ | s | based on diffusive scaling |
| $\tau_{\text{BGK}}$ | relaxation time | 0.8 | – | minimising flow rate error |
| $\eta_0$ | plasma viscosity | 1 | mPa s | same as liquid water |
| $\eta_{\text{cytosol}}$ | cytosol viscosity | 1 | mPa s | assuming $\eta_{\text{cytosol}} = \eta_0$ |
| $R_{\text{rbc}}$ | RBC radius | 4 | $\mu\text{m}$ | present RBC model |
| $T_{\text{rbc}}$ | RBC thickness | 2.5 | $\mu\text{m}$ | present RBC model |
| $A_{\text{rbc}}$ | RBC surface area | 140 | $\mu\text{m}^2$ | present RBC model |
| $V_{\text{rbc}}$ | RBC volume | 100 | $\mu\text{m}^3$ | present RBC model |
| $N$ | number of facets | 2000 | – | $N = 20(r_{\text{rbc}}/\Delta x)^2$ |
| $\Gamma$ | Föppl-von Kármán number | 400 | – | $\Gamma = \kappa_s r_{\text{rbc}}^2 / \kappa_b$ |
| $\tilde{\kappa}_s$ | strain modulus | $1.67 \times 10^{-3}$ | – | resisting in-plane shearing |
| $\tilde{\kappa}_b$ | bending modulus | $4.18 \times 10^{-4}$ | – | resisting off-plane bending |
| $\tilde{\kappa}_\alpha$ | dilation modulus | 0.5 | – | conserving local area |
| $\tilde{\kappa}_A$ | surface modulus | 1 | – | conserving cell surface |
| $\tilde{\kappa}_V$ | volume modulus | 1 | – | conserving cell volume |
| $\tilde{\kappa}_{cc}$ | repulsion intensity | $\tilde{\kappa}_b$ | – | cell-cell interaction |
| $\tilde{\kappa}_{cw}$ | repulsion intensity | $10\tilde{\kappa}_b$ | – | cell-wall interaction |

### S2 Empirical model for blood viscosity in tube flow

Capitalising on various sources of experimental data collected from literature, Pries and co-workers [31] derived an empirical model that depicts the variation of the apparent viscosity of blood *η*_app_ (relative to that of plasma *η*_0_) in cylindrical tubes against the tube diameter *D* (formulated in *μ*m) and discharge haematocrit *H*_D_:

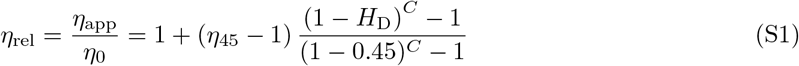

where *η*_45_ is the relative apparent viscosity of blood at a discharge haematocrit *H*_D_ = 0.45 formulated as

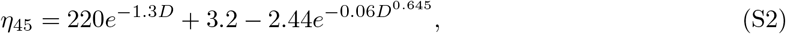

and *C* is a coefficient related to *D* in the form of

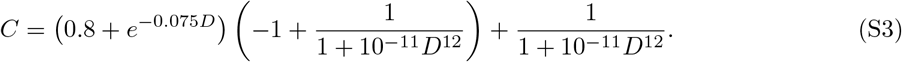

### S3 Spatiotemporal dynamics of the RBC suspension flow

**Figure S1:**
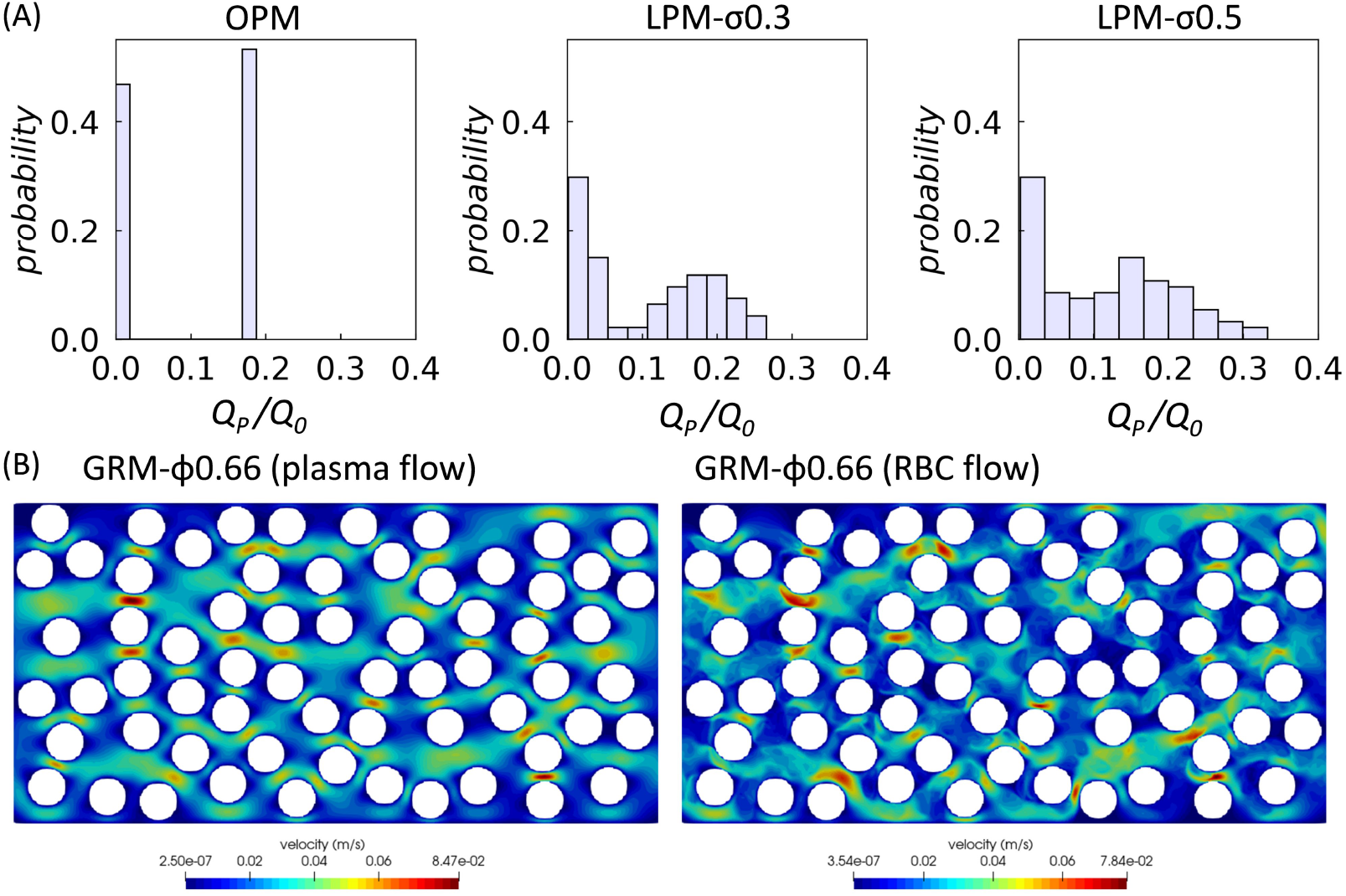
Effect of RBCs interacting with structural disorder on flow distribution within the porous media. (A) Flow rate distributions for steady plasma flows through LPM with increasing levels of disorder *σ* = 0, 0.3, 0.5 (where *σ* = 0 refers to the ordered porous medium), corresponding to the RBC flows *Q*_s_ (*H*_F_ = 20%) in Fig. 4B. Porosity is *ϕ* = 0.57 in all cases. The histograms show flow rates *Q*_p_ evaluated at all throats within the pore space network. *Q*_0_ = 0.4 *μ*L/min is the imposed flow rate at the inlet. (B) Flow channelisation patterns for (left) steady plasma flow and (right) RBC flow (*H*_F_ = 20%) in GRM-*ϕ*0.66 under the same inflow *Q*_0_ = 0.4 *μ*L/min.

**Figure S2:**
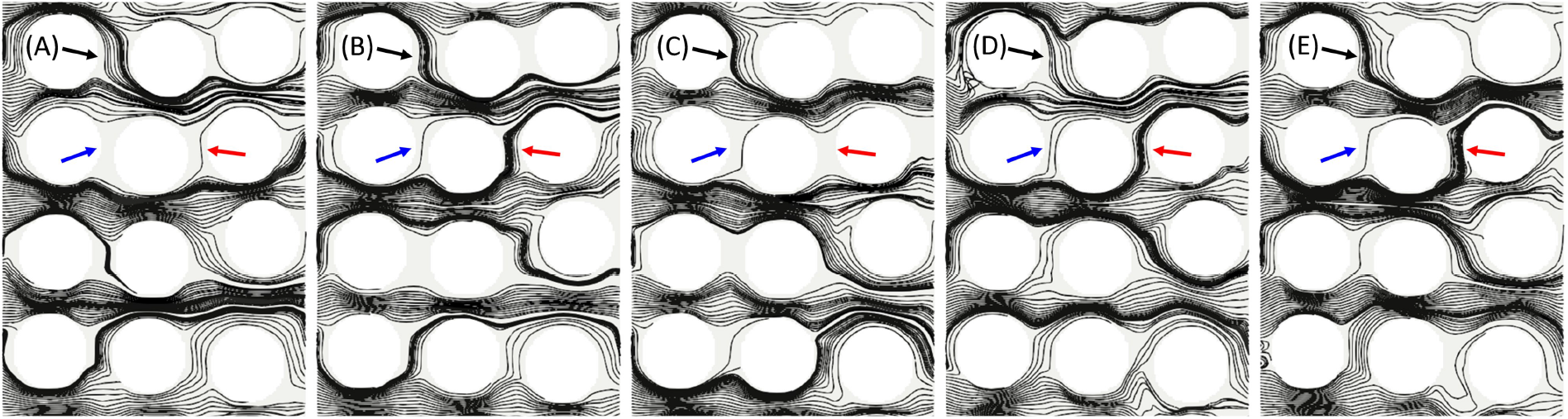
Instantaneous streamlines of the simulation in Fig. 4A. The corresponding time instants are *t* = 134.8 ms, *t* = 137.8 ms, *t* = 140.8 ms, *t* = 143.7 ms, *t* = 146.7 ms, respectively. The temporal variation of the streamlines is caused by the dynamics of the discrete RBCs. The coloured arrows in (A-E) indicate the three throats of interest (“1”, “2”, “3”) marked in panel Fig. 4A.

### S4 Effect of medium porosity and disorder on RBC transport

**Figure S3:**
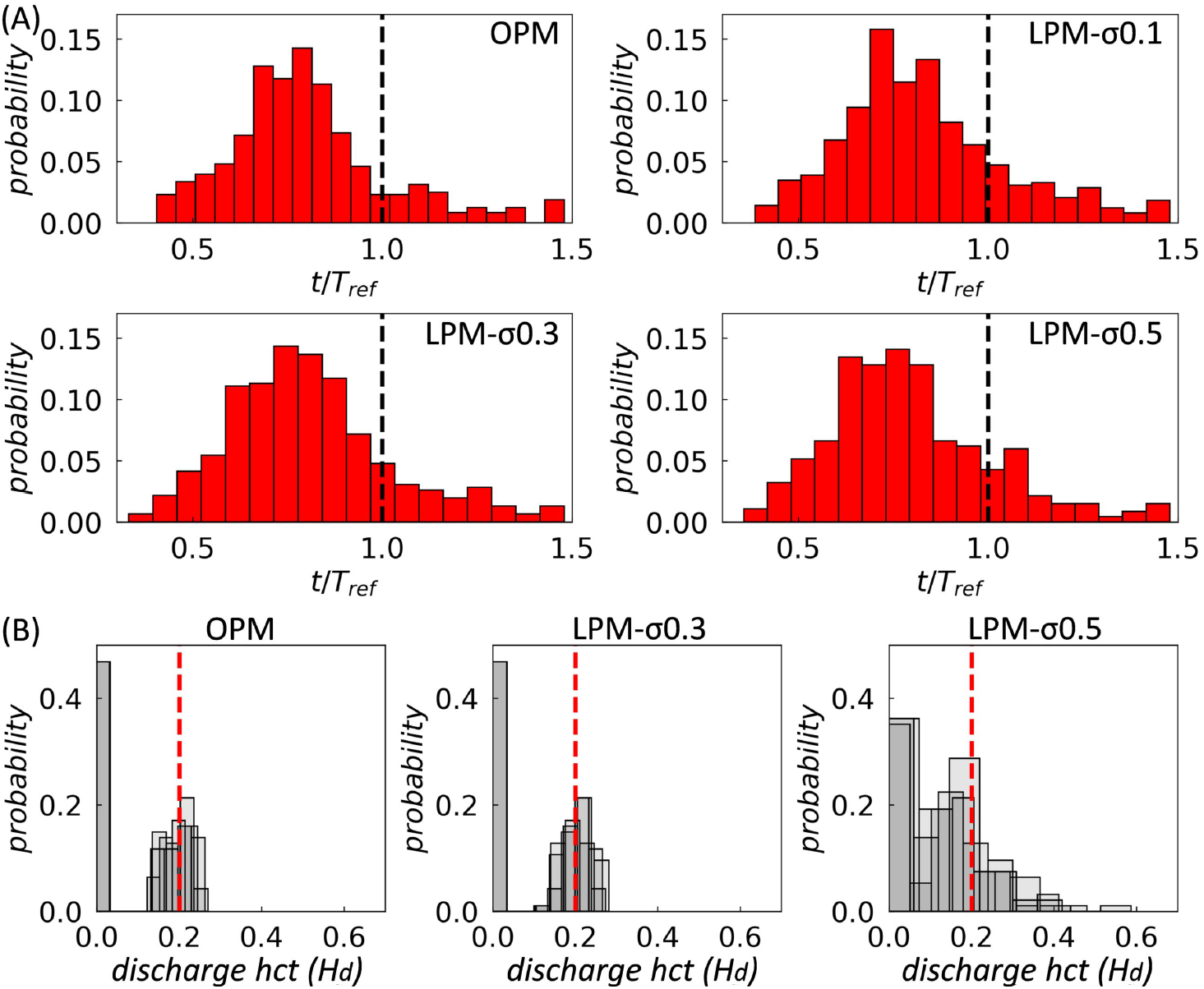
Quantification of particle transit time and discharge haematocrit in other geometries (*ϕ* = 0.57). (A) Transit time distributions in the OPM and LPM geometries (*σ* = 0.1, 0.3, 0.5). (B) Discharge haematocrit distributions in the OPM and weakly disordered LPM geometries (*σ* = 0.1, 0.3).

### S5 Supplementary movies

**Movie1** Video clip of the RBC simulation in OPM-*ϕ*0.57 (*H*_F_ = 0.2).

**Movie2** Video clip of the RBC simulation in LPM-*σ*0.5-*ϕ*0.57 (*H*_F_ = 0.2).

**Movie3** Video clip of the droplet experiment in LPM-*σ*0.5-*ϕ*0.57 (*H*_F_ = 0.2).

**Movie4** Video clip of the RBC simulation in LPM-*σ*0.5-ϕ0.67 (*H*_F_ = 0.2).

**Movie5** Video clip of the RBC simulation in LPM-*σ*0.7-ϕ0.67 (*H*_F_ = 0.2).

**Movie6** Video clip of the RBC simulation in GRM-*ϕ*0.66 (*H*_F_ = 0.2).

